# Follicle-stimulating hormone and luteinizing hormone increase Ca^2+^ in the granulosa cells of mouse ovarian follicles^1^

**DOI:** 10.1101/520122

**Authors:** Jeremy R. Egbert, Paul G. Fahey, Jacob Reimer, Corie M. Owen, Alexei V. Evsikov, Viacheslav O. Nikolaev, Oliver Griesbeck, Russell S. Ray, Andreas S. Tolias, Laurinda A. Jaffe

## Abstract

In mammalian ovarian follicles, follicle stimulating hormone (FSH) and luteinizing hormone (LH) signal primarily through the G-protein G_s_ to elevate cAMP, but both of these hormones can also elevate Ca^2+^ under some conditions. Here we investigate FSH- and LH-induced Ca^2+^ signaling in intact follicles of mice expressing genetically encoded Ca^2+^ sensors, Twitch-2B and GCaMP6s. At a physiological concentration (1 nM), FSH elevates Ca^2+^ within the granulosa cells of preantral and antral follicles. The Ca^2+^ rise begins several minutes after FSH application, peaks at ~10 minutes, remains above baseline for another ~10 minutes, and depends on extracellular Ca^2+^. However, suppression of the FSH-induced Ca^2+^ increase by reducing extracellular Ca^2+^ does not inhibit FSH-induced phosphorylation of MAP kinase, estradiol production, or the acquisition of LH responsiveness. Like FSH, LH also increases Ca^2+^, when applied to preovulatory follicles. At a physiological concentration (10 nM), LH elicits Ca^2+^ oscillations in a subset of cells in the outer mural granulosa layer. These oscillations continue for at least 6 hours and depend on the activity of G_q_ family G-proteins. Suppression of the oscillations by G_q_ inhibition does not inhibit meiotic resumption, but does delay the time to 50% ovulation by about 3 hours. In summary, both FSH and LH increase Ca^2+^ in the granulosa cells of intact follicles, but the functions of these Ca^2+^ rises are only starting to be identified.

**Summary sentence:** Both FSH and LH increase Ca^2+^ in the granulosa cells of intact ovarian follicles from mice expressing genetically encoded sensors.

## INTRODUCTION

In mammals, the G-protein coupled receptors for follicle stimulating hormone (FSH) and luteinizing hormone (LH) mediate events that lead to ovulation of fertilizable eggs. FSH receptors are expressed in granulosa cells of all follicles between the primary and preovulatory stages [1] and mediate the action of FSH to induce granulosa cell proliferation and differentiation, steroidogenesis, and LH receptor expression [2]. LH then acts on its receptors on the mural granulosa cells of preovulatory follicles, leading to oocyte maturation and ovulation [2,3].

Both FSH and LH stimulate the G_s_/cAMP/protein kinase A (PKA) pathway [2], and pharmacological activation of this pathway is, with a few exceptions, sufficient to fully mimic responses to FSH [1,2,4] and LH [2,5–7]. Correspondingly, many of the FSH- and LH-stimulated responses are inhibited by inhibition of PKA [1,2], although non-specificity of some commonly used PKA inhibitors [8] leaves open the question of whether FSH and LH signaling through non-PKA pathways may also be important. In addition to stimulating cAMP production, FSH increases intracellular Ca^2+^ in isolated porcine granulosa cells [9,10], but not in a rat granulosa cell line [11]. LH signaling elevates Ca^2+^ in isolated porcine granulosa cells [12], and in luteinized human granulosa cells [13,14], although not in isolated mouse granulosa cells [15].

These reports raise questions about whether FSH and LH also elevate Ca^2+^ in other species, and in intact follicles, and what functions such Ca^2+^ rises might have in follicular development. Previous studies have suggested possible roles for Ca^2+^ in the FSH-stimulated MAP kinase activation that contributes to estrogen synthesis [16,17], and in LH-stimulated meiotic resumption [18,19]. We chose to investigate these questions using intact follicles, because although many granulosa cell signaling pathways are similar in isolated cells and in intact follicles [2], some are not. For example, in intact follicles, LH causes a decrease in cyclic GMP to a few percent of the basal level [6,20], whereas in isolated granulosa cells, this decrease fails to occur [21] or occurs only partially [22].

Although AM-ester-based Ca^2+^ indicators have been useful for investigating Ca^2+^ dynamics in isolated granulosa cells [9,10,12] and isolated oocytes [23,24], their poor permeation into multilayer tissues precludes their use in isolated follicles. Mice expressing an early generation genetically encoded sensor, TN-XXL [25], had insufficient sensitivity to be useful (our unpublished results), but recently, mice expressing high-affinity optical sensors for Ca^2+^ have been developed [26,27]. Using mouse lines expressing the sensors Twitch-2B and GCaMP6s, we show that FSH and LH both elevate Ca^2+^ in the granulosa cells of intact follicles. We also begin to investigate the physiological functions of the FSH- and LH-induced Ca^2+^ rises.

## MATERIALS AND METHODS

### Mice

Mice globally expressing the Twitch-2B Ca^2+^ sensor [28] were generated at Baylor College of Medicine, using CRISPR/CAS9 technology to target a Cre recombinase-responsive conditional Twitch-2B expression cassette containing the CAG promoter into the Rosa 26 locus. CRISPR/CAS9 and donor sequences were delivered by pronuclear injection into fertilized donor oocytes and implanted into pseudopregnant females. The mice were authenticated by isolating DNA from the founder and from F1 pups, followed by PCR and Southern blot using primers spanning the 3' or 5' insert junctions. To generate globally expressing Twitch-2B mice, conditional mice were bred to a *Bact*-Cre line to permanently delete the LoxP flanked stop cassette and establish a globally expressing sublineage. The mouse line was maintained on a C57BL/6J background.

Measurements were made using homozygotes expressing two copies of the Twitch-2B transgene or with heterozygotes expressing one copy. The concentration of Twitch-2B in the homozygote follicle cytoplasm was estimated to be ~20 µM, based on western blotting (Supplementary Fig. S1A, see methods below), and an estimated cytoplasmic volume per follicle of 20 nl [29]. The mice had normal appearance and were fertile, and follicle-enclosed oocytes from these mice showed a normal time course of nuclear envelope breakdown (NEBD) and ovulated in response to LH (Supplementary Fig. S1B), indicating that the sensor did not perturb physiological function.

Mice that express the GCaMP6s Ca^2+^ sensor Cre-dependently by way of a floxed stop cassette

[27] were obtained from The Jackson Laboratory (Bar Harbor, ME; stock number 024106; B6;129S6/J strain). To induce global GCaMP6s expression, these mice were bred with *Hprt*-Cre mice [30] that were originally obtained from The Jackson Laboratory (stock number 004302), and that had been backcrossed onto a C57BL/6J background. The resulting line was maintained on a C57BL/6J background. Measurements were made using mice heterozygous for GCaMP6s. These mice had normal appearance and were fertile.

Genotyping of both Ca^2+^ sensor lines was accomplished by observing the fluorescence of ears and tails using goggles fitted with FHS/EF-3GY1 emission filters (BLS Ltd, Budapest, Hungary). Heterozygotes and homozygotes were distinguished by PCR genotyping for the wildtype allele using primers listed in Supplementary Table S1.

Wildtype C57BL/6J mice were used for MAPK western blots, estradiol measurements, and measurements of nuclear envelope breakdown and ovulation. For most of these experiments, the wildtype mice were purchased from The Jackson Laboratory; a few wildtype mice were from the breeding colonies for the lines expressing Ca^2+^ sensors. All experiments with mice were performed at the University of Connecticut Health Center, and all animal protocols were approved by the University of Connecticut Health Center Animal Care Committee.

### Isolation and culture of follicles

Follicles of the indicated sizes were dissected from prepubertal (24-27 day old) mice, and cultured on Millicell organotypic membranes (Merck Millipore, Cork, Ireland; PICMORG50) in MEMα medium (12000-022, Invitrogen) with 25 mM NaHCO_3_, 75 µg/ml penicillin G, 50 µg/ml streptomycin, a mixture of 5 µg/ml insulin, 5 µg/ml transferrin, and 5 ng/ml selenium (Sigma-Aldrich, St. Louis, MO, #I1884), and 3 mg/ml bovine serum albumin (MP Biomedicals, #103700). Where indicated, 1 nM FSH was included in the medium. To reduce Ca^2+^ in the medium, EGTA was added, and the resulting free Ca^2+^ concentrations were calculated using MaxChelator (https://somapp.ucdmc.ucdavis.edu/pharmacology/bers/maxchelator/downloads.htm) [31]).

Follicles were used for experiments 24-30 hours after isolation. Highly purified ovine FSH (AFP7558C) and ovine LH (oLH-26) were obtained from A.F. Parlow (National Hormone and Peptide Program, Torrance, CA). The follicles, which were spheres when dissected, flattened to disks after culture on the Millicell membrane, thus facilitating imaging. Diameters described in the text refer to measurements at the time of dissection, and do not include the residual theca layer. For determining meiotic resumption in response to LH, follicle-enclosed oocytes were checked hourly for the loss of the nucleolus and nuclear envelope.

### Confocal Imaging

For confocal microscopy, follicles were placed in a 100 µl drop of medium in the channel of a “µ-slide” that connects two ports (ibidi GmbH, Martinsried, Germany, #80176 custom ordered without adhesive; see [6]). Slides with a channel height of 100 µm (ibidi custom order) were used for follicles 140-250 µm in diameter; 290-360 µm follicles were placed in a 200 µm-deep slide. A thin layer of silicone grease was applied to the outer edge of the slide and a glass coverslip (ibidi # 10812) was placed on the silicone to gently immobilize the follicle in the channel. FSH, LH, or control medium was applied by perfusing 200 μl of the solution through the channel by way of the ports. For recordings at 1-6 hours after LH treatment, LH was applied to the follicles on Millicell membranes; just before imaging, follicles were transferred to a µ-slide in medium containing LH.

Follicles were imaged on a Zeiss LSM 5 Pascal confocal microscope using a C-Apochromat 40X/1.2 numerical aperture objective with Immersol W immersion medium (Carl Zeiss Microscopy). Twitch-2B CFP was excited using a 440 nm laser (Toptica Photonics, Victor, NY) using a power range of 1-5%. The dichroic (510DLCP) and CFP and YFP emission filters (HQ480/40M and HQ535/50M, respectively) were from Chroma Technology (Rockingham, VT). GCaMP6s was imaged using a 488 nm laser and an HQ525/50M emission filter. A warm air blower (Nevtek ASI-400, Burnsville, VA) was used to maintain a stage temperature of 32-35 °C. For Twitch-2B, the pinhole was fully open (13 µm optical section); for GCaMP6s recordings, a 5 µm optical section was used. 12-bit scans at 512 x 512 resolution were collected every 10 seconds for up to 50 minutes. Background correction was applied by subtracting the averaged autofluorescence of 3-5 wildtype follicles imaged under identical conditions. Because ~28% of the light collected by the YFP emission filter is actually emitted by CFP, we subtracted 28% of the background-subtracted YFP intensity before calculating the CFP/YFP ratio.

### RNA sequencing data analysis

Transcriptome data of mural granulosa cells from 2-month-old C57BL/6J mice [32] in FASTQ format were downloaded from the European Nucleotide Archive (ENA; https://www.ebi.ac.uk/ena). Sequence alignments were performed using the RNA STAR tool [33] within the Galaxy platform [34]. Reads were aligned to the mouse reference genome (GRCm38, a.k.a. mm10). Gene model for splice junctions, in GTF file format, was obtained from ENSEMBL ftp site (release 92). The GTF file was filtered to contain only entries with the “gene_biotype” value set to “protein_coding” in the “attributes” field. To calculate gene expression, we counted the numbers of reads aligned to exons using featureCounts [35] version 1.6.0.6 from within the Galaxy platform, with the value of “minimum bases of overlap” set to 30. To account for the depth of sequencing among datasets, we normalized count expression data to counts per million (CPM) [36] by using the formula *CPM*_*i*_ = 10^6^ × *C*_*i*_ / *T*_*a*_, where *CPM*_*i*_ is the CPM value for the gene *i*, *C*_*i*_ is the number of counts reported by featureCounts, and *T*_*a*_ is the total number of reads in the sample aligned to the genome. To identify all genes encoding Ca^2+^ channels, gene symbols for all mouse genes annotated to “GO:0015085: calcium ion transmembrane transporter activity” were downloaded from Mouse Genome Informatics [37] (130 genes, downloaded on 09/05/2018). This list was edited to include only those genes encoding plasma membrane proteins that transport Ca^2+^. Ca^2+^ channels in organelles, Ca^2+^ channel regulatory proteins, and TRP channels that are not significantly permeable to Ca^2+^ [38] were removed from the list.

### Western blotting

Western blots for quantifying the Twitch-2B concentration were probed with a primary antibody against GFP (Cell Signaling Technology, Beverly, MA; #2555); this antibody also recognizes CFP and YFP. The Epac2-camps protein that was used as a standard for these blots has a similar CFP and YFP-containing structure as Twitch-2B, and >90% purity [39]. Western blots for detecting MAP kinase phosphorylation were probed with primary antibodies against phospho-MAPK1/3 (Thr202/Tyr204) and total MAPK1/3 (Cell Signaling Technology, #4370 and #4696, respectively). MAPK1 is also known as ERK2 or p42MAPK, and MAPK3 is also known as ERK1 or p44MAPK. The antibody used to detect PDE1A was from Proteintech (Rosemont, IL, #12442-2-AP). For all western blots except Fig. S4, antibody binding was detected using fluorescent secondary antibodies (LI-COR Biosciences, Lincoln, NE; IRDye800CW and IRDye680RD) and a LICOR Odyssey imaging system. The secondary antibody for Fig. S4 was conjugated to horseradish peroxidase (Santa Cruz Biotechnology, Dallas, TX) and detected with ECL Prime (GE Healthcare, Chicago, IL) and a CCD camera (G:BOX Chemi XT4, Syngene, Frederick, MD). Further antibody details are included in Supplementary Table S2. Signal intensities were measured using ImageJ (https://imagej.nih.gov/ij/).

### Sources of reagents

YM-254890 was obtained from Wako Chemicals (Richmond, VA). EGTA, cadmium chloride, nickel chloride, and nifedipine were from Sigma-Aldrich. ELISA kits for measurement of 17β-estradiol were from Cayman Chemical (Ann Arbor, MI).

### Statistics

Analyses were conducted as described in the figure legends using Prism 6 (GraphPad Software, Inc, La Jolla, CA). All values indicate mean ± standard error of the mean (SEM). Multiple t-tests (either with or without ANOVA) were corrected for multiple comparisons using the Holm-Sidak correction when comparing all groups to each other, or the Dunnett correction when comparing all groups to a single control.

## RESULTS AND DISCUSSION

### Measurement of an FSH-induced increase in Ca^2+^ in the granulosa cells of intact ovarian follicles expressing the Twitch-2B FRET sensor

Using Twitch-2B-expressing follicles ranging in diameter from ~140 µm (preantral) to ~320 µm (fully grown antral), we measured Ca^2+^ levels before and after addition of FSH. The follicles were imaged by confocal microscopy, with the focus on the oocyte equator (Fig. 1A). Twitch-2B fluorescence was seen in the granulosa cells, but was too low to be useful for measurements in the oocyte. In some of the smaller follicles, the fluorescence was uniform throughout the granulosa cells (Fig. 1A), although in others, it was dimmer in the interior granulosa layers. In larger follicles, the fluorescence was consistently dimmer in the interior layers (see Fig. 4A). For this reason, measurements were made from the outer 25 µm region of granulosa cell layer, except as indicated. Twitch-2B was also present in the residual theca/blood vessel layer surrounding the follicle, and was expressed at a higher level than in the granulosa cells. The amount of adhering theca was variable (compare Figs 1A and 4A). Because the theca layer was not intact, we did not investigate FSH-induced Ca^2+^ changes in this region.

**Figure 1.**
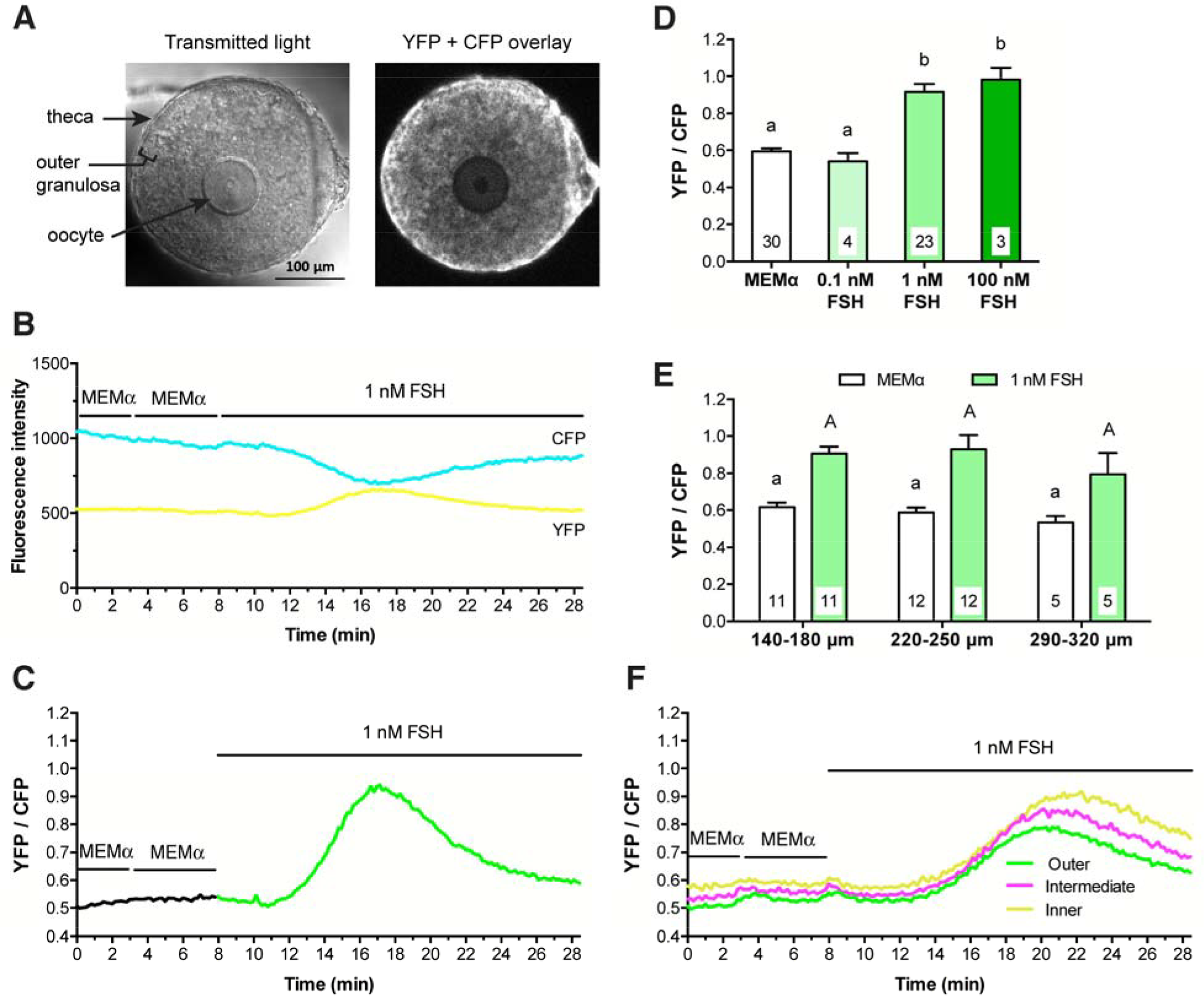
FSH increases intracellular Ca^2+^ in the mural granulosa cells of intact follicles, as detected by Twitch-2B. **(A)** Representative transmitted light and CFP + YFP fluorescence images of a follicle after a 28-hour culture on a Millicell membrane. Prior to flattening on the membrane, the follicle measured ~180 µm in diameter. Note that the residual theca cells that surround the follicle and that were not removed by dissection show higher fluorescence intensity than the granulosa cells, indicating the presence of more Twitch-2B protein in the theca cells. **(B)** Representative traces showing changes in YFP and CFP fluorescence before and after perfusion with control medium (MEMα) or 1 nM FSH. Following FSH perfusion, the two channels change in opposite directions, indicating an increase in FRET (same follicle as A). **(C)** Representative trace showing the YFP/CFP ratio before and after treatment with 1 nM FSH (same follicle as B). The increase in YFP/CFP ratio indicates that 1 nM FSH induces a transient Ca^2+^ increase in the granulosa cells. **(D)** Both 1 nM and 100 nM FSH induce Ca^2+^ increases of similar magnitude. Peak YFP/CFP ratios during perfusion of control medium (MEMα) and the indicated concentrations of FSH. Follicle diameters at the time of isolation from the ovary were 140-250 µm. **(E)** The Ca^2+^ response to 1 nM FSH is similar in follicles of different sizes. Peak YFP/CFP ratios before and after perfusion of 1 nM FSH, for follicles of the indicated diameters (those in the 140-180 µm and 220-250 µm groups are the same follicles as in A). Numbers within the bars indicate the number of follicles tested. Different letters indicate significant differences (p < 0.05) after one-way ANOVA (D) or two-way ANOVA (E), followed by t-tests with the Holm-Sidak correction for multiple comparisons. For (E), lowercase letters indicate comparisons among MEMα groups; uppercase letters reflect comparisons between groups treated with FSH. All values represent mean ± s.e.m. **(F)** The FSH-induced Ca^2+^ rise is similar in different regions of preantral follicles. Traces show fluorescence intensity as a function of time for 3 concentric regions of a representative follicle: in the outer 25 µm (green), the middle 25-50 µm (magenta), and the inner 25 µm (yellow). The follicle used for these measurements was 160 µm in diameter at the time of isolation. Similar records were obtained from 6 other follicles with diameters ranging from 140 to 220 µm.

Binding of Ca^2+^ to Twitch-2B increases FRET between CFP and YFP, such that an increase in the YFP/CFP emission ratio measured after CFP excitation indicates an increase in free Ca^2+^; the EC_50_ of Twitch-2B is ~200 nM [28]. In response to perfusion of 1 nM FSH, YFP emission in the granulosa cells increased and CFP emission decreased (Fig. 1B,C), indicating an increase in Ca^2+^. The Ca^2+^ rise began several minutes after FSH application. Ca^2+^ levels reached a peak at about 10 minutes after the initial FSH exposure, and remained above baseline for at least another 10 minutes.

### The FSH-induced Ca^2+^ increase occurs at a physiological concentration of FSH, in follicles of different sizes, and for preantral follicles, throughout the entire granulosa cell compartment

In Twitch-2B-expressing follicles, a concentration of 1 nM FSH was sufficient to elevate Ca^2+^ to an almost maximal level, while no rise in Ca^2+^ was seen with 0.1 nM FSH (Fig. 1D). Follicles of all size classes tested responded similarly to 1 nM FSH (Fig. 1E). The 1 nM concentration of FSH required to stimulate a rise in Ca^2+^ correlates well with the concentration of FSH needed to optimally stimulate follicular development in our culture system. In this system, 0.3 nM FSH causes ~80% of follicles to acquire the ability to resume meiosis in response to LH, whereas 1 nM FSH causes 100% of follicles to become LH responsive [7]. During the reproductive cycle, the FSH concentration in mouse serum reaches a peak of ~1 nM [40]. Thus, the concentration of FSH that we find to be required to elevate Ca^2+^ in the granulosa cells of isolated follicles is in a physiologically relevant range.

The measurements described above were made in the outer 25 µm of the granulosa cell layer, due to low expression of the Ca^2+^ sensor in the follicle interior. However, for some of the smaller follicles (7 of the 23 in the 140-250 µm diameter range, in Fig. 1E), Twitch-2B fluorescence in the inner regions of the follicle was sufficient to measure. In these follicles, the FSH-induced Ca^2+^ rise had an approximately similar amplitude and time course in all regions of the granulosa cell compartment (Fig. 1F).

### Lack of Ca^2+^ buffering by the Twitch-2B sensor

Expression of the Twitch-2B sensor could conceivably buffer cytosolic Ca^2+^, reducing the signal amplitude. We assessed this possibility by comparing mice heterozygous or homozygous for Twitch-2B. CFP fluorescence was about two-fold higher in homozygous follicles (Supplementary Fig. S2A), reflective of Twitch-2B protein expression levels. However, the baseline YFP/CFP ratio and the peak ratio in response to FSH were not different between heterozygous and homozygous follicles (Supplementary Fig. S2B), indicating that Twitch-2B expression did not alter basal Ca^2+^ levels or attenuate the FSH-induced Ca^2+^ increase. Thus the ~20 µM cytosolic concentration of the Twitch-2B sensor in homozygous follicles (see Materials and Methods) is not a significant buffer, consistent with previous determinations that ~100 µM of the Ca^2+^ buffer BAPTA is required to blunt Ca^2+^ rises in other cells [41,42].

### The FSH-induced Ca^2+^ increase occurs uniformly in all cells in the outer granulosa cell region, as detected in follicles expressing the GCaMP6s sensor

To test whether Ca^2+^ increases uniformly in all cells in the outer 25 µm granulosa region, we imaged the Ca^2+^ increase using follicles from mice expressing the GCaMP6s sensor [27], which shows increased fluorescence when Ca^2+^ is bound; the EC_50_ for GCaMP6s is ~140 nM [43]. GCaMP6s is not a ratiometric indicator, so is less useful for comparing different follicles, but it has the advantage of allowing direct visualization of the spatial distribution of Ca^2+^ within a single follicle. As seen with Twitch-2B, GCaMP6s fluorescence was fairly uniform in the granulosa cells of smaller follicles, although undetectable in the oocyte (Fig. 2A), but was restricted to the outer mural granulosa cells in larger follicles (see Figs 5A and 6A below).

**Figure 2.**
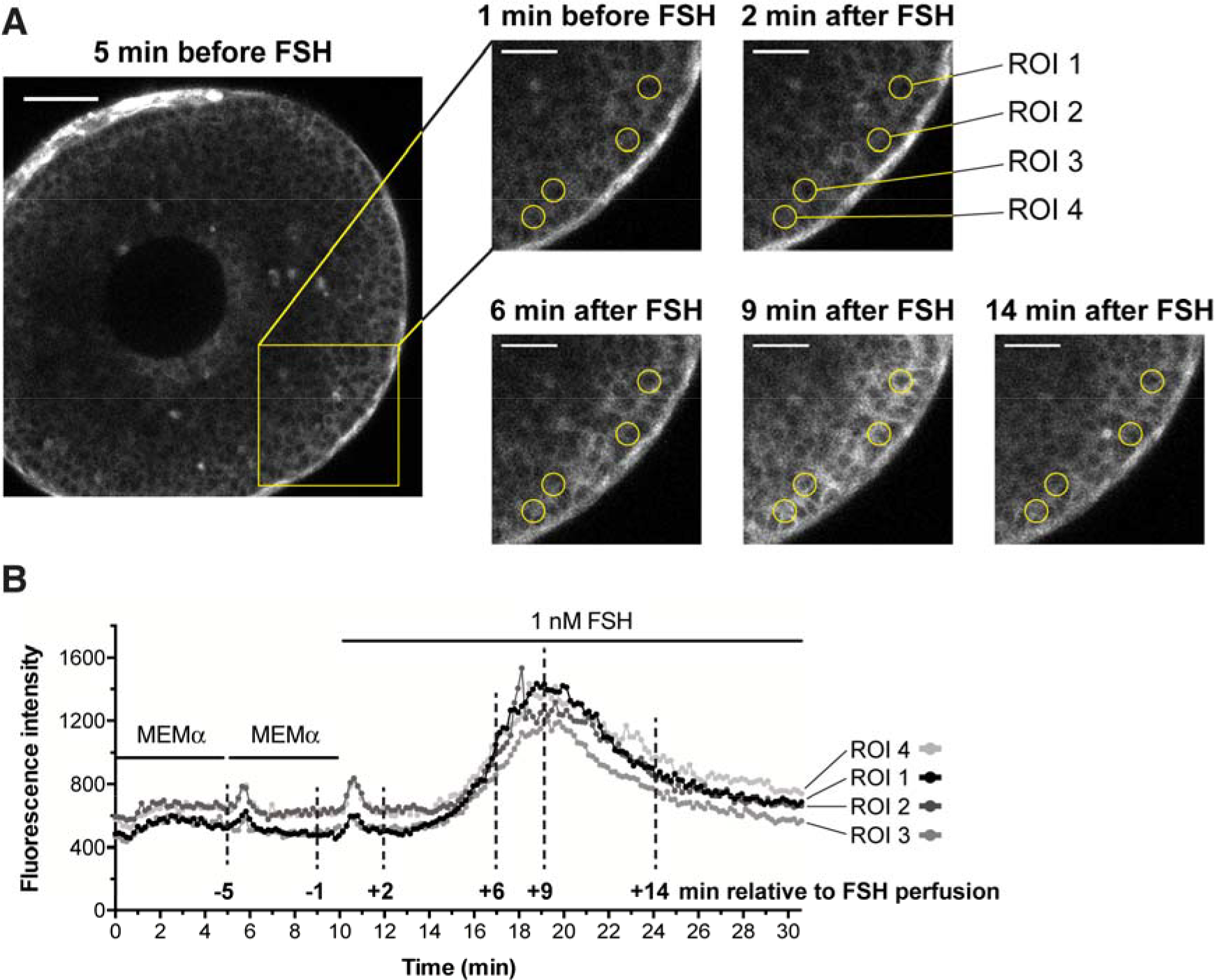
FSH increases Ca^2+^ uniformly in all outer granulosa cells, as detected with GCaMP6s. **(A)** A small antral follicle from a mouse expressing the GCaMP6s Ca^2+^ sensor (~220 µm-diameter at the time of isolation 24 hours previously). GCaMP6s fluorescence intensity increases with increasing Ca^2+^ levels, allowing Ca^2+^ rises in individual cells to be visualized. The yellow box shows the area of enlargements at different time points before and after FSH perfusion; the time points corresponding to these images are marked on the traces in (B) with vertical dashed lines. Yellow circles highlight the regions of interest (ROIs) (10 µm diameter), positioned approximately over single cells, that were measured to generate the traces in (B). Note that after FSH, the entire outer granulosa layer increases in brightness to a similar extent. Scale bar in full image is 50 µm; scale bars in cropped images represent 25 µm. Supplementary movie S1 shows the entire time series from which these images were selected. **(B)** GCaMP6s fluorescence intensity (relative Ca^2+^ levels) for each of the 4 ROIs labeled above. Note that the granulosa cells show a small Ca^2+^ transient in response to the mechanical stimulus during perfusion, independent of FSH. Representative of 6 follicles.

Following perfusion with 1 nM FSH, Ca^2+^ increased uniformly throughout the outer granulosa cell region, and all cells had similar Ca^2+^ dynamics (Fig. 2A,B and supplementary movie 1). The kinetics of the Ca^2+^ increase and subsequent decrease were similar to those observed with Twitch-2B (compare Figs 1C and 2B).

Like Twitch-2B, GCaMP6s was also expressed in the theca layer, and its fluorescence there was brighter than in the granulosa cells (see Figs 2A, 5A, 6A). It is unknown if the higher fluorescence in the theca was due to higher expression of the sensor protein or to higher basal Ca^2+^.

### The FSH-induced Ca^2+^ increase is due to influx from the extracellular solution

To investigate if the FSH-induced Ca^2+^ increase is due to Ca^2+^ influx from the extracellular solution, we added 2.0 mM EGTA to the medium, which contained 1.8 mM CaCl_2_, thus lowering extracellular free Ca^2+^ to ~0.002 mM (see Materials and Methods). After 20 minutes, we perfused FSH onto the follicle while measuring the Twitch-2B signal. Under these conditions, FSH did not cause a detectable increase in Ca^2+^ in the granulosa cells (Fig. 3A,B). Likewise, when 1.6 mM EGTA was added to the medium, to reduce extracellular Ca^2+^ to 0.2 mM, FSH did not cause a detectable Ca^2+^ increase (Fig. 3A,B). These results indicate that the FSH-induced Ca^2+^ increase results from Ca^2+^ influx from the extracellular solution. Consistent with this conclusion, exposure to 10 μM YM-254890, which inhibits signaling by G_q_-family G-proteins [44,45; see Figs 4E and 6A,B below] and thus inhibits Ca^2+^ release from the endoplasmic reticulum, did not prevent the Ca^2+^ rise in response to FSH (Fig. 3A,B).

**Figure 3.**
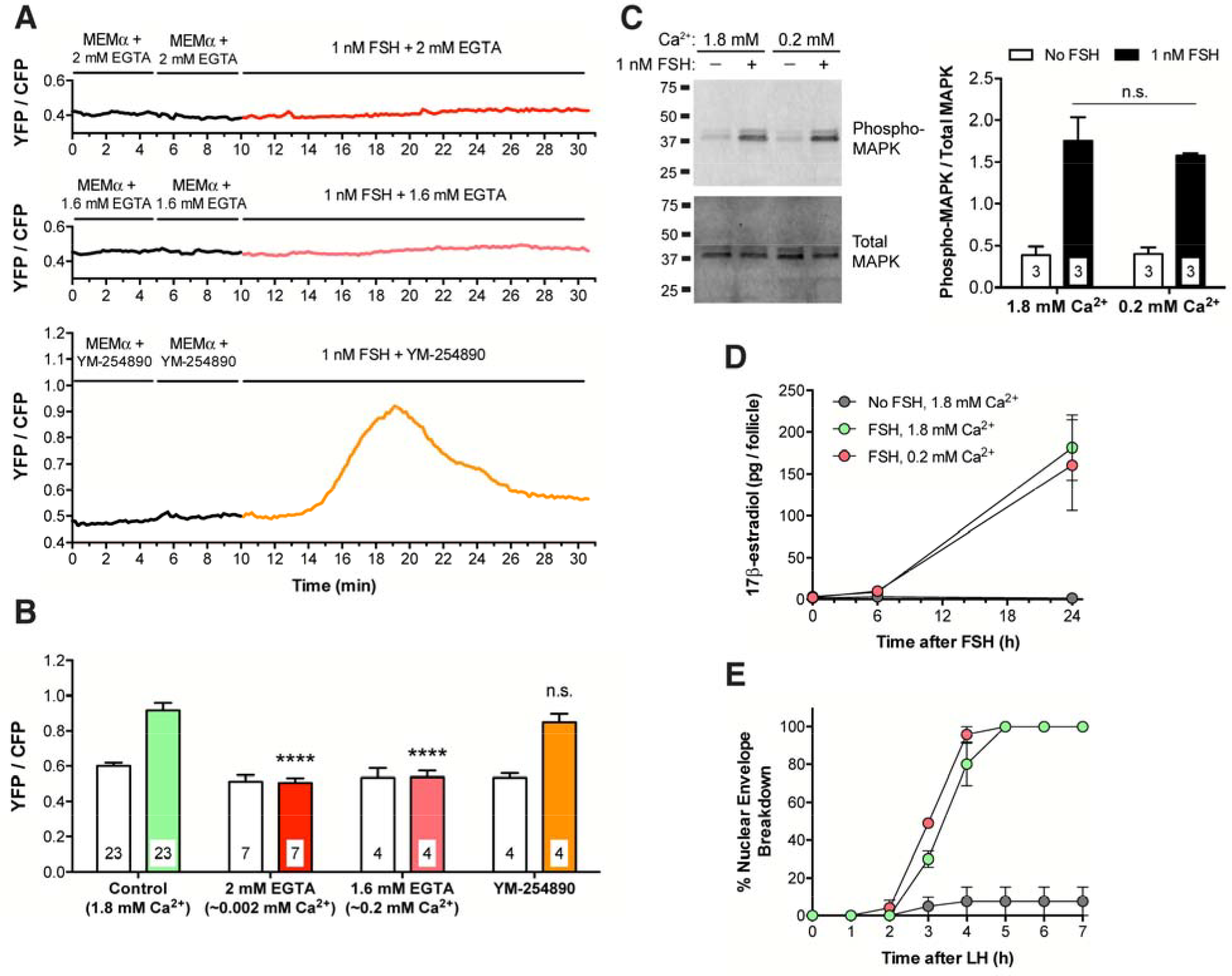
The FSH-induced Ca^2+^ increase in granulosa cells of intact follicles requires extracellular Ca^2+^, but MAPK phosphorylation, 17β-estradiol production, and LH-induced nuclear envelope breakdown occur independently of the Ca^2+^ increase. **(A)** Representative traces showing no FSH-induced Ca^2+^ increase in follicles (220-250 µm diameter) in media with low extracellular Ca^2+^, and no effect of the G_q_-family G-protein inhibitor YM-254890 (10 µM) on the FSH-induced Ca^2+^ increase. MEMα, which contains 1.8 mM CaCl_2_, was mixed with either 2 mM EGTA (~0.002 mM free Ca^2+^; upper panel), or 1.6 mM EGTA (~0.2 mM free Ca^2+^; middle panel). Follicles were pre-incubated for ~20 min in these EGTA solutions, or for 60 min in YM-254890 (lower panel), before addition of FSH. **(B)** Peak YFP/CFP ratios before and after perfusion of 1 nM FSH in the presence of varying extracellular Ca^2+^ concentrations, buffered by EGTA addition as in A, or in the presence of YM-254890 (10 µM). The ability of YM-254890 to inhibit LH-induced Ca^2+^ elevation (see Figs 4E and 6A,B below) served as a positive control for the permeability of this inhibitor. Open bars indicate the peak YFP/CFP ratio after MEMα perfusion; filled bars indicate peak YFP/CFP ratio following FSH perfusion. Following 2-way ANOVA, the peak ratios following FSH perfusion in each group were compared to the control (green) bar by t-tests with the Dunnett correction for multiple comparisons; **** indicates p < 0.0001, n.s. indicates p > 0.05. None of the peak ratios measured during MEMα perfusion (open bars) were significantly different from the control (all p > 0.05). **(C)** FSH-induced MAPK phosphorylation occurs independently of extracellular Ca^2+^. Left: Representative western blot for phospho- and total MAPK in lysates of 180-250 µm follicles incubated for 20-30 min in medium with either 1.8 mM or 0.2 mM Ca^2+^ (prepared as in A), and then for 20 min in these same media with or without 1 nM FSH. Right: Ratio of phospho-/total MAPK from 3 independent experiments; n.s. indicates p > 0.05 by unpaired t-test. **(D)** FSH-induced estradiol production occurs independently of extracellular Ca^2+^. 290-360 µm follicles were incubated with 1 nM FSH in media containing either 1.8 mM or 0.2 mM Ca^2+^. Medium samples were collected at the indicated times for estradiol analysis. Each point represents 3 experiments using 14-20 follicles for each treatment. **(E)** FSH-induced acquisition of LH responsiveness occurs independently of extracellular Ca^2+^. 290-360 µm follicles were incubated for 24 hours with 1 nM FSH in media with either 1.8 mM or 0.2 mM Ca^2+^; 10 nM LH was then added to each dish and the time course of nuclear envelope breakdown was determined. Each point represents 3 experiments using 12-19 follicles for each condition. Same legend as in (D). Values in (B-E) represent mean ± s.e.m.

**Figure 4:**
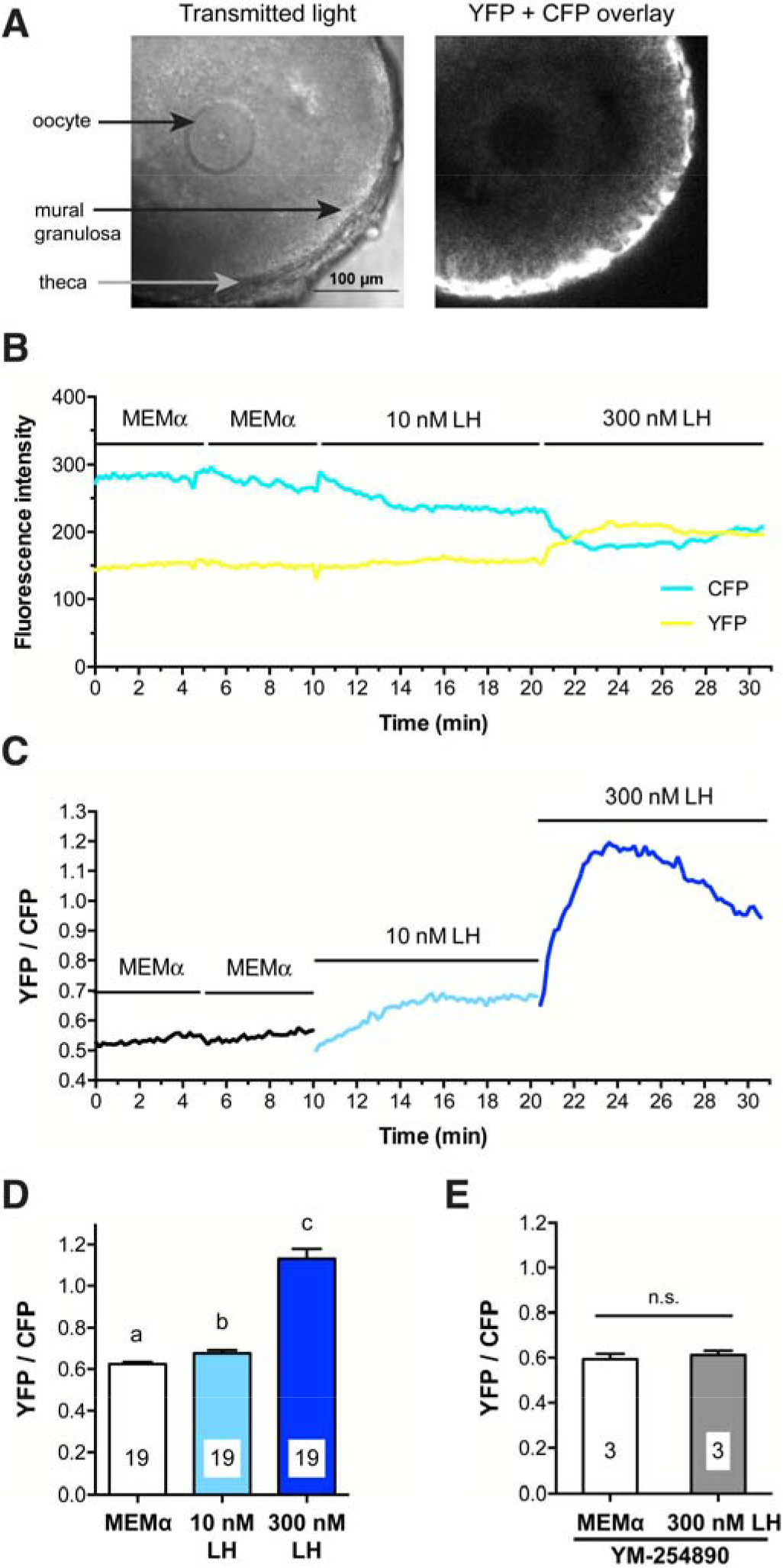
LH causes a G_q_-family G-protein-dependent increase in mural granulosa Ca^2+^, as detected with Twitch-2B. **(A)** Transmitted light and CFP + YFP fluorescence images of a large antral follicle after 25-hour culture on a Millicell membrane in the presence of 1 nM FSH. Prior to flattening on the membrane, the follicle measured ~320 µm in diameter. Note that the theca cells express more Twitch-2B than the granulosa cells. **(B)** YFP and CFP fluorescence for the follicle shown in (A) during sequential perfusions of MEMα, 10 nM LH, and 300 nM LH. **(C)** YFP/CFP ratio during sequential perfusions of MEMα,10 nM LH, and 300 nM LH, for the follicle shown in B. **(D)** Peak YFP/CFP ratios before and after perfusion of 10 nM and 300 nM LH. Different letters indicate significant differences (p < 0.001) after repeated measures ANOVA with the Holm-Sidak correction for multiple comparisons. **(E)** Peak YFP/CFP ratios before and after perfusion of 300 nM LH, following inhibition of G_q_-family G-proteins by a 1-2.5 hour pretreatment with 10 µM YM-254890; n.s. indicates p > 0.05 by paired t-test. Values in (D,E) represent mean ± s.e.m.

**Figure 5.**
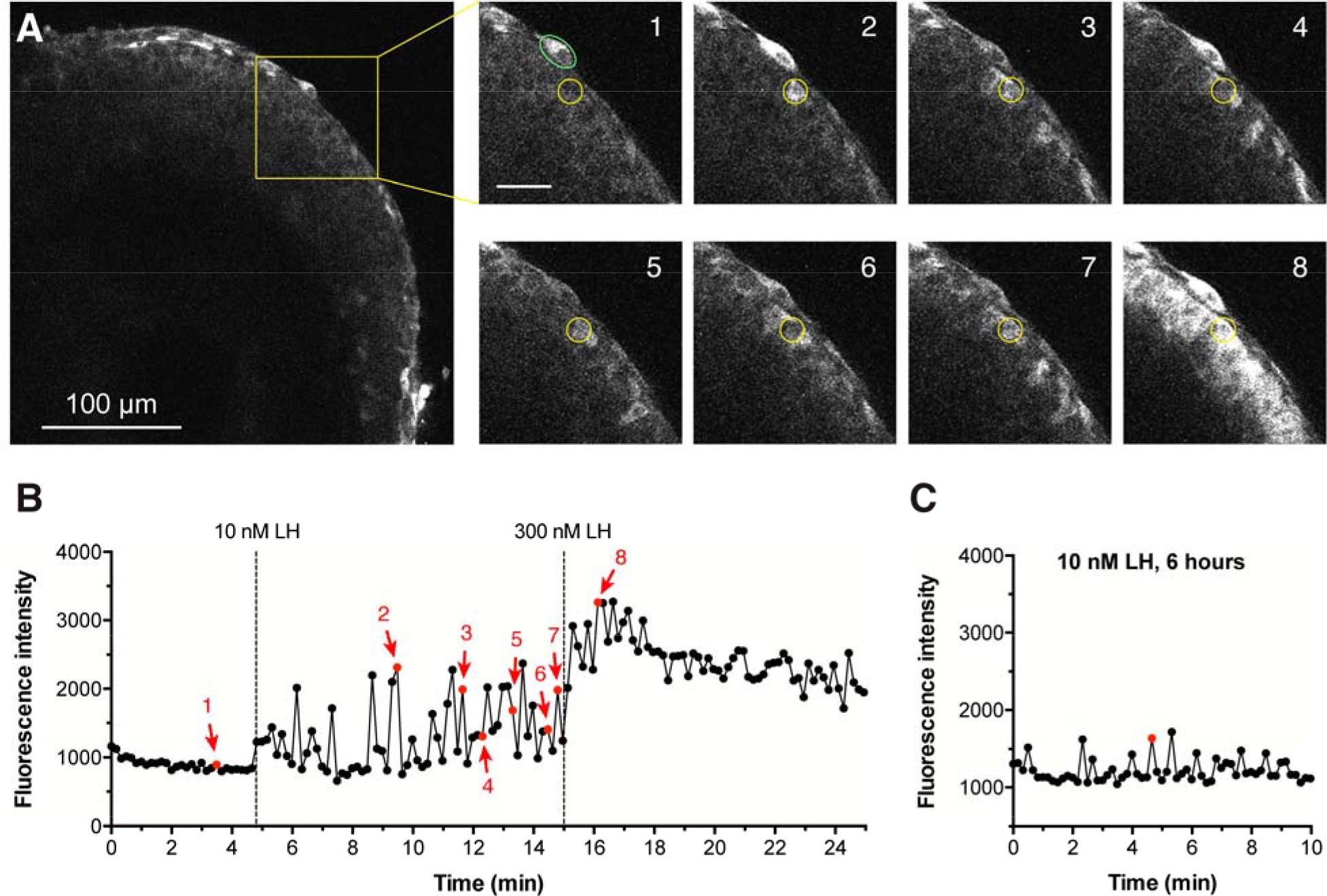
LH induces Ca^2+^ oscillations in mural granulosa cells, as detected with GCaMP6s. **(A)** The left hand image shows a large antral follicle after 25-hour culture in the presence of 1 nM FSH. The yellow box outlines an area for which enlarged images are shown at 8 time points: before LH (image 1), after perfusion of 10 nM LH (images 2-7), and after perfusion of 300 nM LH (image 8); scale bar = 25 µm. The yellow circle (10 µm in diameter) indicates a mural granulosa cell that undergoes Ca^2+^ oscillations in response to 10 nM LH. Calcium oscillations are also visible in several other mural granulosa cells. The green oval indicates a theca cell that undergoes calcium responses in response to 10 nM LH. Supplementary movie S2 shows the entire time series from which these images were selected, and supplementary movie S3 shows another example. **(B)** Fluorescence intensity before and after LH perfusion, for the granulosa cell in (A) that is indicated by the yellow circle. Images were taken at 10-sec intervals. Points in red correspond to the numbered images in (A). Representative of 4 follicles. **(C)** Ca^2+^ oscillations in the outer mural granulosa cells continue for at least 6 hours after treatment with 10 nM LH. The graph shows the fluorescence intensity as a function of time for one such cell. Representative of 2 follicles each at 1, 2, and 4 hours, and 3 follicles at 6 hours after LH. Supplementary movie S4 shows an example at 2 hours after LH, and supplementary movie S5 shows the full time series from which data shown in C were measured.

To identify candidate ion channels potentially mediating the FSH-induced Ca^2+^ influx, we analyzed gene expression data from next generation sequencing of mouse mural granulosa cell mRNA [32] (Supplementary Table S3). Among the genes encoding plasma membrane Ca^2+^ permeable channels, the most highly expressed was *Trpm7*, followed by 2 voltage-gated Ca^2+^ channels: *Cacna1h* (T type) and *Cacna1a* (P/Q type). Other plasma membrane Ca^2+^ permeable channels, including a voltage-gated L type channel, a glutamate receptor, a purinergic receptor, and pannexin 1 were expressed at somewhat lower levels. In an attempt to inhibit some of these channels, we tested several voltage-sensitive Ca^2+^ channel blockers, Cd^2+^, Ni^2+^, and nifedipine [46]. At the concentrations tested, none inhibited the FSH-induced Ca^2+^ increase (Supplementary Fig. S3), which is not surprising because multiple channel types were detected in RNAseq transcriptome data of mural granulosa cells (Supplementary Table S3), likely obscuring suppression of Ca^2+^ influx. The possible role of TRPM7 channels, for which specific inhibitors are not yet available [47], remains to be investigated, as does the question of how activation of the FSH receptor results in opening of these or other Ca^2+^ channels.

### The function of the FSH-induced Ca^2+^ increase is unknown

Previous studies of isolated rat granulosa cells have indicated that FSH-induced phosphorylation of the mitogen-activated protein kinases MAPK1 and MAPK3 depends on extracellular Ca^2+^ [16]. MAPK1/3 phosphorylation is of particular interest because it is essential for the FSH-induced increase in synthesis of aromatase, which catalyzes the synthesis of estradiol [17]. Therefore, we investigated the effect of reducing extracellular Ca^2+^ on the FSH-induced phosphorylation of MAPK1/3 and synthesis of estradiol in mouse follicles. However, suppression of the FSH-induced Ca^2+^ rise by addition of 1.6 mM EGTA to the medium did not inhibit FSH-induced MAPK1/3 phosphorylation (Fig. 3C) or estradiol synthesis (Fig. 3D). It also did not prevent the FSH-induced acquisition of the ability to resume meiosis in response to LH (Fig. 3E). Our findings indicate that the FSH-induced Ca^2+^ rise is not required for several biological functions. Whether the Ca^2+^ rise is part of a back-up system for these functions, or whether it regulates other functions such as FSH-induced granulosa cell proliferation or suppression of apoptosis, remains to be investigated.

### LH causes persistent Ca^2+^ oscillations in the granulosa cells of intact ovarian follicles

To investigate whether LH increases Ca^2+^ in the granulosa cells of intact follicles, we isolated follicles with diameters of 290-360 µm, and cultured them for 24-30 hours in the presence of 1 nM FSH to induce expression of LH receptors. In follicles expressing Twitch-2B, the fluorescence signal was strong in the outer mural granulosa cells, but in the inner mural granulosa cells, cumulus cells, and oocyte, the signal was too weak to be useful (Fig. 4A). Therefore, measurements were made from the outer 25 µm layer of granulosa cells. Perfusion of 10 nM LH caused a barely detectable Ca^2+^ increase, as indicated by a small increase in the YFP/CFP emission ratio of Twitch-2B (Fig. 4B,C,D). In response to subsequent perfusion of 300 nM LH, Ca^2+^ began to increase immediately, reached a peak at about 5 minutes, and remained above baseline for at least another 5 minutes (Fig. 4B,C,D). The peak concentration of LH in the serum during the mouse reproductive cycle is about 1 nM [48,49], but 10 nM is the minimum concentration of ovine LH needed to cause meiotic resumption in all follicles under our experimental conditions [7], perhaps related to use of LH from a different species.

We next tested whether the Ca^2+^ elevation in response to 10 nM LH might be more evident using GCaMP6s, which has an EC_50_ of ~140 nM vs ~200 nM for Twitch-2B. GCaMP6s is also more sensitive because Ca^2+^ causes a proportionately larger change in fluorescence intensity than that seen with Twitch-2B (compare Figs 1B and 2B, and Figs 4B and 5B below). When a follicle expressing GCaMP6s was exposed to 10 nM LH, some individual cells in the outermost mural granulosa layer showed oscillating rises in Ca^2+^, beginning within a minute after perfusion and occurring once or twice per minute throughout the 10 minute recording period (Fig. 5A,B; supplementary movies S2 and S3). During each oscillation, Ca^2+^ remained above baseline for ~10-30 seconds. When the follicle was subsequently perfused with 300 nM LH, Ca^2+^ elevation occurred throughout the 25 μm outer region of the mural granulosa cells (Fig. 5A,B).

To investigate how long the Ca^2+^ oscillations in response to LH continued, we applied 10 nM LH to GCaMP6s-expressing follicles on a Millicell membrane, and then 1, 2, 4, or 6 hours later, transferred them to an imaging chamber. Ca^2+^ oscillations occurred for at least 6 hours after LH application (Fig. 5C; Supplementary Movies S4 and S5), though the amplitude relative to the baseline appeared to be reduced when compared to oscillations immediately after LH (see Fig. 5B). As noted above, the isolated follicles used for these studies had a variable number of theca cells adhering to their surfaces. Although we did not systematically investigate LH-induced responses in the theca, we consistently observed oscillations in GCaMP6s fluorescence intensity when LH was perfused (Figure 5A and supplementary movies S2 and S3). Further investigation of LH-induced calcium oscillations in the theca cells should be done using preparations of ovarian tissue that retain an intact theca layer (for example, see ref. [50]).

### The LH-induced Ca^2+^ increases require the activity of a G_q_-family G-protein

The LH-induced Ca^2+^ elevations in the granulosa cells were blocked by 10 μM of the G_q_-family G-protein inhibitor YM-254890 (Figs 4E and 6A,B; supplementary movie S6). 10 µM YM-254890 inhibits signaling by G_q_-family G-proteins (G_q_, G_11_, and G_14_), but not by other G-proteins (G_s_, G_i_, G_o_, G_13_), or by voltage-gated or ATP-gated Ca^2+^ channels [44,45]. These data indicate that, unlike the FSH-induced Ca^2+^ rise, the LH-induced Ca^2+^ oscillations are most likely initiated by IP_3_-induced Ca^2+^ release from the endoplasmic reticulum. Consistently, previous studies have shown that LH induces IP_3_ production in isolated granulosa cells of rat [22] and mouse [51].

### Inhibition of G_q_-family G-proteins does not inhibit LH-induced meiotic resumption, but does delay ovulation

Previous studies have suggested that an LH-induced Ca^2+^ rise in the granulosa cells might contribute to causing meiotic resumption [18,19]. However, inhibition of the LH-induced Ca^2+^ oscillations with the G_q_-family inhibitor YM-254890 did not inhibit or delay meiotic resumption in mouse follicle-enclosed oocytes (Fig. 6C). This result confirms a previous finding that G_q_ family G-proteins are not required for meiotic resumption [51]. However, since the Ca^2+^-activated cGMP phosphodiesterase PDE1 is expressed in mouse granulosa cells (Egbert et al., 2016, and Supplementary Fig. S4), the LH-induced Ca^2+^ rise would increase cGMP phosphodiesterase activity, and thus would contribute to decreasing cGMP. Therefore, while the Ca^2+^ rise is not required for LH-induced meiotic resumption, it could be a component of a “failsafe” system that ensures that LH signaling lowers cGMP such that meiosis proceeds.

**Figure 6.**
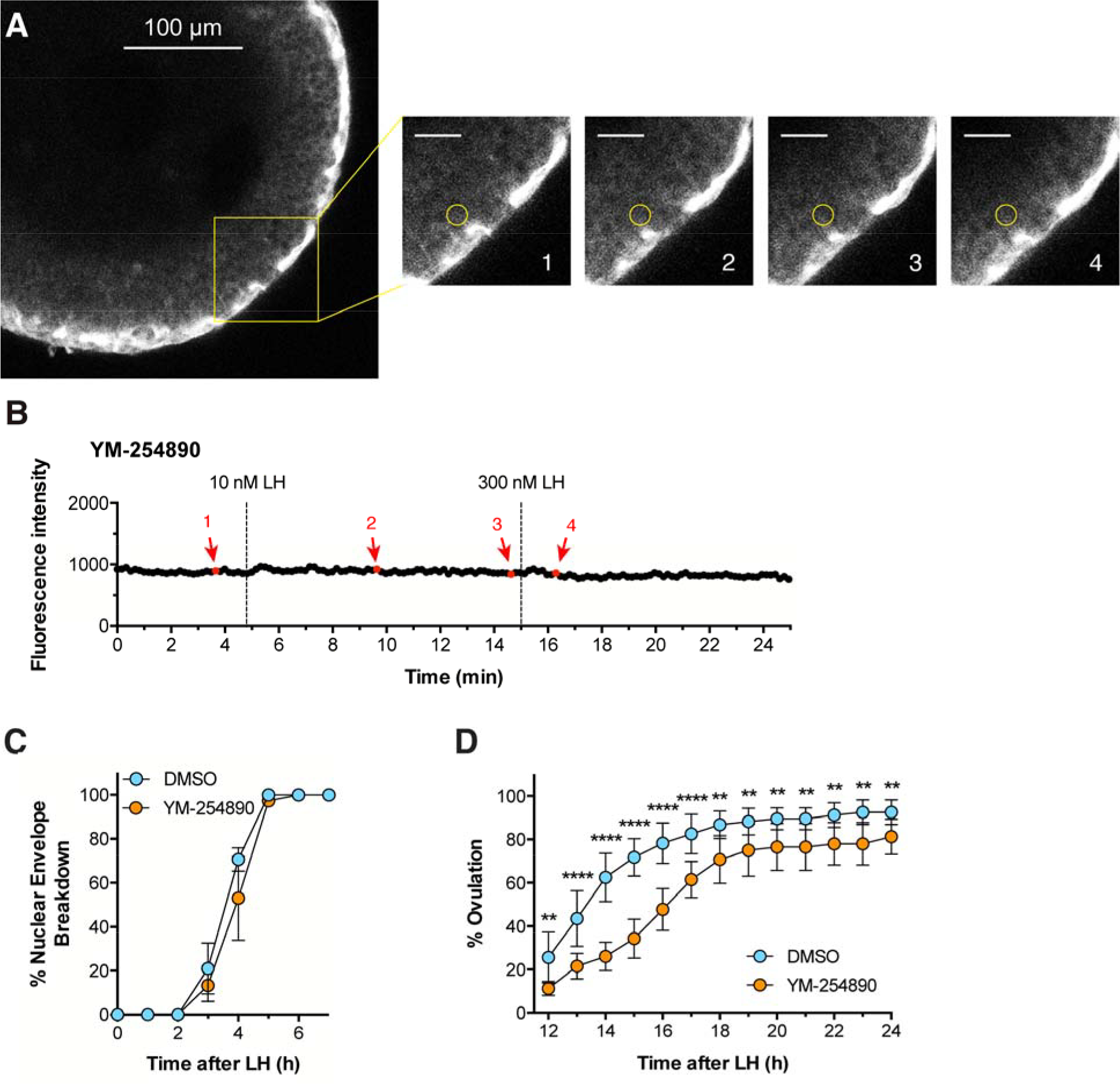
The G_q_-family G-protein inhibitor YM-254890 prevents Ca^2+^ oscillations in response to LH, and delays ovulation. Experiments were performed using follicles 290-360 µm in diameter that had been cultured for ~24 hours in the presence of 1 nM FSH. **(A,B)** A follicle expressing GCaMP6s was pre-incubated with 10 µM YM-254890 for 60 min before adding 10 nM LH. The yellow box outlines an area for which enlarged images are shown at 4 time points: before LH (image 1), after perfusion of 10 nM LH (images 2 and 3), and after perfusion of 300 nM LH (image 4); scale bar = 25 µm. The yellow circle (10 µm in diameter) indicates a mural granulosa cell for which the fluorescence intensity is graphed in B. Supplementary movie S6 shows the entire time series from which these images were selected. Representative of 3 follicles.**(C,D)** Inhibition of G_q_-family G-proteins by preincubation of follicles with 10 µM YM-254890 for 60 min before addition of 10 nM LH does not inhibit nuclear envelope breakdown, but does delay ovulation. Graphs in C and D show the mean ± s.e.m of 3 and 4 experiments, respectively, with 10-18 follicles each. Data in (D) were analyzed by repeated measures 2-way ANOVA, with the Holm-Sidak correction for multiple comparisons applied to the t-tests between the DMSO and YM-254890 groups at each time point; ** indicates p < 0.01 and **** indicates p < 0.0001.

Inhibition of G_q_ family G-proteins with YM-254890 did have an inhibitory effect on LH-induced ovulation (Fig. 6D). 50% of control follicles had ovulated by ~13 hours after LH application, whereas in the presence of 10 µM YM-254890, 50% ovulation did not occur until ~16 hours. These results with isolated follicles confirm the inhibitory effect on ovulation that was seen in mice in which G_q_ and G_11_ expression in the granulosa cells was inactivated genetically [51]. Thus, the LH-induced Ca^2+^ oscillations may signal together with cAMP to stimulate ovulation. Other possible functions of LH-induced Ca^2+^ oscillations, in both the granulosa and theca cells, remain to be investigated.

## ACKNOWLEDGEMENTS

We thank Rachael Norris and Leia Shuhaibar for their participation in initiating this project using earlier Ca^2+^ measurement methods, Siu-Pok Yee and Deborah Kaback for their expertise and advice on mouse genome modification and colony maintenance, and Tracy Uliasz, Giulia Vigone, Mary Hunzicker-Dunn, Aylin Hanyaloglu, Lixia Yue, and Paul Brehm for useful discussions. The Tolias lab thanks Zheng Huan Tan for colony management.

## REFERENCES

[1] Hardy K, Fenwick M, Mora J, Laird M, Thomson K, Franks S. Onset and heterogeneity of responsiveness to FSH in mouse preantral follicles in culture. Endocrinology 2017; 158:134–147.

[2] Hunzicker-Dunn M, Mayo K. Gonadotropin signaling in the ovary. In Knobil and Neill’s Physiology of Reproduction, 4th Edition. T.M. Plant, and A.J. Zeleznik, editors. Academic Press/San Diego, 2015; pp. 895–945.

[3] Jaffe LA, Egbert JR. Regulation of mammalian oocyte meiosis by intercellular communication within the ovarian follicle. Ann Rev Physiol 2017; 79:237–260.

[4] Puri P, Little-Ihrig L, Chandran U, Law NC, Hunzicker-Dunn M, Zeleznik AJ. Protein kinase A: a master kinase of granulosa cell differentiation. Sci Rep 2016; 6:28132.

[5] Reizel Y, Elbaz J, Dekel N. Sustained activity of the EGF receptor is an absolute requisite for LH-induced oocyte maturation and cumulus expansion. Mol Endocrinol 2010; 24:402–411.

[6] Shuhaibar LC, Egbert JR, Norris RP, Lampe PD, Nikolaev VO, Thunemann M, Wen L, Feil R, Jaffe LA. Intercellular signaling via cyclic GMP diffusion through gap junctions in the mouse ovarian follicle. Proc Natl Acad Sci USA 2015; 112:5527–5532.

[7] Vigone G, Shuhaibar LC, Egbert JR, Uliasz TF, Movsesian MA, Jaffe LA. Multiple cAMP phosphodiesterases act together to prevent premature oocyte meiosis and ovulation. Endocrinology 2018; 159:2142–2152.

[8] Davies SP, Reddy H, Caivano M, Cohen P. Specificity and mechanism of action of some commonly used protein kinase inhibitors. Biochem J 2000; 351:95–105.

[9] Flores JA, Veldhuis JD, Leong DA. Follicle-stimulating hormone evokes an increase in intracellular free calcium ion concentrations in single ovarian (granulosa) cells. Endocrinology 1990; 127:3172–3179.

[10] Flores JA, Leong DA, Veldhuis JD. Is the calcium signal induced by follicle-stimulating hormone in swine granulosa cells mediated by adenosine cyclic 3’, 5’-monophosphate-dependent protein kinase? Endocrinology 1992; 130:1862–1866.

[11] Grieshaber NA, Boitano S, Ji I, Mather JP, Ji TH. Differentiation of granulosa cell line: Follicle-stimulating hormone induces formation of lamellipodia and filopodia via the adenylyl cyclase/cyclic adenosine monophosphate signal. Endocrinology 2000; 141:3461–3470.

[12] Flores JA, Aguirre C, Sharma OP, Veldhuis JD. Luteinizing hormone (LH) stimulates both intracellular calcium ion ([Ca^2+^]_i_) mobilization and transmembrane cation influx in single ovarian (granulosa) cells: Recruitment as a cellular mechanism of LH-[Ca^2+^]_i_ dose response. Endocrinology 1998; 139:3606–3612.

[13] Lee PSN, Buchan AMJ, Hsueh AJW, Yuen BH, Leung PCK. Intracellular calcium mobilization in response to the activation of human wild-type and chimeric gonadotropin receptors. Endocrinology 2002; 143:1732–1740.

[14] Jonas KC, Chen S, Virta M, Mora J, Franks S, Huhtaniemi I, Hanyaloglu AC. Temporal reprogramming of calcium signaling via crosstalk of gonadotropin receptors that associate as functionally asymmetric heteromers. Sci Rep 2018; 8:2239.

[15] Webb RJ, Bains H, Cruttwell C, Carroll J. Gap-junctional communication in mouse cumulus-oocyte complexes: implications for the mechanism of meiotic maturation. Reproduction 2002; 123:41–52.

[16] Cottom J, Salvador LM, Maizels ET, Reierstad S, Park Y, Carr DW, Davare MA, Hell JW, Palmer SS, Dent P, Kawakatsu H, Ogata M, Hunzicker-Dunn M. Follicle-stimulating hormone activates extracellular signal-regulated kinase but not extracellular signal-regulated kinase kinase through a 100-kDa phosphotryosine phosphatase. J Biol Chem 2003; 278:7167–7179.

[17] Donabauer EM, Hunzicker-Dunn ME. Extracellular signal-regulated kinase (ERK)-dependent phosphorylation of Y-box-binding protein 1 (YB-1) enhances gene expression in granulosa cells in response to follicle-stimulating hormone (FSH). J Biol Chem 2016; 291:12145–12160.

[18] Goren S, Oron Y, Dekel N. Rat oocyte maturation: role of calcium in hormone action. Mol Cell Endocrinol 1990; 72:131–138.

[19] Egbert JR, Uliasz TF, Shuhaibar LC, Geerts A, Wunder F, Kleiman RJ, Humphrey JM, Lampe PD, Artemyev NO, Rybalkin SD, Beavo JA, Movsesian MA, Jaffe LA. Luteinizing hormone causes phosphorylation and activation of the cyclic GMP phosphodiesterase PDE5 in rat ovarian follicles, contributing, together with PDE1 activity, to the resumption of meiosis. Biol Reprod 2016; 94(5):110,1–11.

[20] Egbert JR, Shuhaibar LC, Edmund AB, Van Helden DA, Robinson JW, Uliasz TF, Baena V, Geerts A, Wunder F, Potter LR, Jaffe LA. Dephosphorylation and inactivation of the NPR2 guanylyl cyclase in the granulosa cells contributes to the LH-induced cGMP decrease that causes resumption of meiosis in rat oocytes. Development 2014; 141:3594–3604.

[21] Lee K-B, Zhang M, Sugiura K, Wigglesworth K, Uliasz T, Jaffe LA, Eppig JJ. Hormonal coordination of natriuretic peptide type C and natriuretic peptide receptor 3 expression in mouse granulosa cells. Biol Reprod 2013; 88:1–9.

[22] Davis JS, Weakland LL, West LA, Farese RV. Luteinizing hormone stimulates the formation of inositol trisphosphate and cyclic AMP in rat granulosa cells. Biochem J 1986; 238:597–604.

[23] Cheon B, Lee H-C, Wakai T, Fissore RA. Ca^2+^ influx and the store-operated Ca^2+^ entry pathway undergo regulation during mouse oocyte maturation. Mol Biol Cell 2013; 24:1396–1410.

[24] Carvacho I, Ardestani G, Lee HC, McGarvey K, Fissore RA, Lykke-Hartmann K. TRPM7-like channels are functionally expressed in oocytes and modulate post-fertilization embryo development in mouse. Sci Rep 2016; 6:34236.

[25] Mank M, Santos AF, Direnberger S, Mrsic-Flogel TD, Hofer SB, Stein V, Hendel T, Reiff DF, Levelt C, Borst A, Bonhoeffer T, Hübener M, Griesbeck O. A genetically encoded calcium indicator for chronic in vivo two-photon imaging. Nat Methods 2008; 5:805–811.

[26] Direnberger S, Mues M, Micale V, Wotjak CT, Dietzel S, Schubert M, Scharr A, Hassan S, Wahl-Schott C, Biel M, Krishnamoorthy G, Griesbeck O. Biocompatibility of a genetically encoded calcium indicator in a transgenic mouse model. Nat Commun 2012; 3:1031.

[27] Madisen L, Garner AR, Shimaoka D, Chuong AS, Klapoetke NC, Li L, van der Bourg A, Niino Y, Egolf L, Monetti C, Gu H, Mills M, Cheng A, Tasic B, Nguyen TN, Sunkin SM, Benucci A, Nagy A, Miyawaki A, Helmchen F, Empson RM, Knöpfel T, Boyden ES, Reid RC, Carandini M, Zeng H. Transgenic mice for intersectional targeting of neural sensors and effectors with high specificity and performance. Neuron 2015; 85:942–958.

[28] Thestrup T, Litzlbauer J, Bartholomäus I, Mues M, Russo L, Dana H, Kovalchuk Y, Liang Y, Kalamakis G, Laukat Y, Becker S, Witte G, Geiger A, Allen T, Rome LC, Chen T-W, Kim DS, Garaschuk O, Griesinger C, Griesbeck O. Optimized ratiometric calcium sensors for functional *in vivo* imaging of neurons and T lymphocytes. Nat Methods 2014; 11:175–182.

[29] Norris RP, Freudzon M, Nikolaev VO, Jaffe LA. Epidermal growth factor receptor kinase activity is required for gap junction closure and for part of the decrease in ovarian follicle cGMP in response to LH. Reproduction 2010; 140:655–662.

[30] Tang S-HE, Silva FJ, Tsark WMK, Mann JR. A Cre/loxP-deleter transgenic line in mouse strain 129S1/SvlmJ. Genesis 2002; 32:199–202.

[31] Bers DM, Patton CW, Nuccitelli R. A practical guide to the preparation of Ca^2+^ buffers. Meth Cell Biol 2010; 99:1–26.

[32] Kawasaki K, Sumitomo J, Shiroguchi K, Endo TA, Sugiura K. (2016). Assessing transcriptomes of mouse cumulus cells, mural granulosa cells, and cumulus-oocyte complexes. https://www.ncbi.nlm.nih.gov/geo/query/acc.cgi?acc=GSE80326.

[33] Dobin A, Davis CA, Schlesinger F, Drenkow J, Zaleski C, Jha S, Batut P, Chaisson M, Gingeras TR. STAR: Ultrafast universal RNA-seq aligner. Bioinformatics 2013; 29:15–21.

[34] Goecks J, Nekrutenko A, Taylor J, Galaxy Team. Galaxy: A comprehensive approach for supporting accessible, reproducible, and transparent computational research in the life sciences. Genome Biol 2010; 11:R86.

[35] Liao Y, Smyth GK, Shi W. Featurecounts: An efficient general purpose program for assigning sequence reads to genomic features. Bioinformatics 2014; 30:923–930.

[36] Mortazavi A, Williams BA, McCue K, Schaeffer L, Wold B. Mapping and quantifying mammalian transcriptomes by RNA-seq. Nat Methods 2008; 5:621–628.

[37] Smith CL, Blake JA, Kadin AJ, Richardson JE, Bult CJ. Mouse Genome Database (MGD)-2018:) knowledgebase for the laboratory mouse. Nucleic Acids Res 2018; 46:D836–D842.

[38] Mulier M, Vriens J, Voets T. TRP channel pores and local calcium signals. Cell Calcium 2017; 66:19–24.

[39] Nikolaev VO, Bunemann M, Hein L, Hannawacker A, Lohse MJ. Novel single chain cAMP sensors for receptor-induced signal propagation. J Biol Chem 2004; 279:37215–37218.

[40] Visser JA, Durlinger ALL, Peters IJJ, van den Heuvel ER, Rose UM, Kramer P, de Jong FH, Themmen APN. Increased oocyte degeneration and follicular atresia during the estrous cycle in anti-Müllerian hormone null mice. Endocrinology 2007; 148:2301–2308.

[41] Kline D. Calcium-dependent events at fertilization of the frog egg: injection of a calcium buffer blocks ion channel opening, exocytosis, and formation of pronuclei. Dev Biol 1988; 126:346–361.

[42] Rozov A, Burnashev N, Sakmann B, Neher E. Transmitter release modulation by intracellular Ca^2+^ buffers in facilitating and depressing nerve terminals of pyramidal cells in layer 2/3 of the rat neocortex indicates a target cell-specific difference in presynaptic calcium dynamics. J Physiol 2001; 531:807–826.

[43] Chen TW, Wardill TJ, Sun Y, Pulver SR, Renninger SL, Baohan A, Schreiter ER, Kerr RA, Orger MB, Jayaraman V, Looger LL, Svoboda K, Kim DS. Ultrasensitive fluorescent proteins for imaging neuronal activity. Nature 2013; 499:295–300.

[44] Takasaki J, Saito T, Taniguchi M, Kawasaki T, Moritani Y, Hayashi K, Kobori M. A Novel Gα_q/11_-selective inhibitor. J Biol Chem 2004; 279:47438–47445.

[45] Nishimura A, Kitano K, Takasaki J, Taniguchi M, Mizuno N, Tago K, Hakoshima T, Itoh H. Structural basis for the specific inhibition of heterotrimeric G_q_ protein by a small molecule. Proc Natl Acad Sci USA 2010; 107:13666–13671.

[46] Catterall WA, Perez-Reyes E, Snutch TP, Striessnig J. International Union of Pharmacology. XLVIII. Nomenclature and structure-function relationships of voltage-gated calcium channels. Pharmacol Rev 2005; 57:411–425.

[47] Chubanov V, Ferioli S, Gudermann T. Assessment of TRPM7 functions by drug-like small molecules. Cell Calcium 2017; 67:166–173.

[48] Rodriguez KF, Couse JF, Jayes FL, Hamilton KJ, Burns KA, Taniguchi F, Korach KS. Insufficient luteinizing hormone-induced intracellular signaling disrupts ovulation in preovulatory follicles lacking estrogen receptor-β Endocrinology 2010; 151:2826–2834.

[49] Czieselsky K, Prescott M, Porteous R, Campos P, Clarkson J, Steyn FJ, Campbell RE, Herbison AE. Pulse and surge profiles of luteinizing hormone secretion in the mouse. Endocrinology 2016; 157:4794–4802.

[50] Komatsu K, Masubuchi S. Observation of the dynamics of follicular development in the ovary. Reprod Med Biol 2017; 16:21–27.

[51] Breen SM, Andric N, Ping T, Xie F, Offermans S, Gossen JA, Ascoli M. Ovulation involves the luteinizing hormone-dependent activation of G_q/11_ in granulosa cells. Mol Endocrinol 2013; 27:1483–1491.

