## Supplementary Data for "Follicle-stimulating hormone and luteinizing hormone increase Ca^2+^ in the granulosa cells of mouse ovarian follicles^1^"

Table S1. PCR primers used for detecting the wild-type allele for genotyping of Twitch-2B and GCaMP6s mice.

| Mouse line | Forward primer, 5'-3' | Reverse primer, 5'-3' | Product size (bp) |
|---|---|---|---|
| **Twitch-2B** | CTCTGCTGCCTCCTGGCTTCTGAG | CTCCGAGGCGGATCACAAGC | 325 |
| **GCaMP6s** | AAGGGAGCTGCAGTGGAGTA | GGATATGAAGTACTGGGCTC | 550 |

Table S2. Antibodies used for western blotting.

| Antibody ID | Target | Vendor | Cat. # | Clonality | Host Organism |
|---|---|---|---|---|---|
| AB_10692764 | Green fluorescent protein (GFP) | Cell Signaling Technology | 2555 | Polyclonal | Rabbit |
| AB_2315112 | MAPK1/3 (p42/44 MAPK), phospho (Thr202 / Tyr 204) | Cell Signaling Technology | 4370 | Monoclonal, clone D13.14.4E | Rabbit |
| AB_390780 | MAPK1/3 (p42/44 MAPK), total | Cell Signaling Technology | 4696 | Monoclonal, clone L34F12 | Mouse |
| AB_2162893 | Phosphodiesterase 1A (PDE1A) | Proteintech Group | 12442-2-AP | Polyclonal | Rabbit |
| AB_2651127 | Rabbit IgG, IRDye 800CW-conjugated | LI-COR Biosciences | 925-32211 | Polyclonal | Goat |
| AB_2651128 | Mouse IgG, IRDye 680RD-conjugated | LI-COR Biosciences | 925-68070 | Polyclonal | Goat |
| AB_631746 | Rabbit IgG, HRP-conjugated | Santa Cruz Biotechnology | sc-2004 | Polyclonal | Goat |

Table S3. $Ca^{2+}$ permeable plasma membrane channels identified by RNA sequencing of mouse mural granulosa cells (average of 2 samples from 2-month old C57BL/6 mice; see Materials and Methods). The right-hand column indicates counts per million (CPM) and standard error of the mean (SEM) for each gene. Genes with average CPM ≥ 25 are highlighted.

| Symbol | Name | CPM (mean ± SEM) |
|---|---|---|
| *Cacna1a* | calcium channel, voltage-dependent, P/Q type, alpha 1A subunit | 47.8±2.2 |
| *Cacna1b* | calcium channel, voltage-dependent, N type, alpha 1B subunit | 0.4±0.0 |
| *Cacna1c* | calcium channel, voltage-dependent, L type, alpha 1C subunit | 3.4±1.2 |
| *Cacna1d* | calcium channel, voltage-dependent, L type, alpha 1D subunit | 28.7±0.4 |
| *Cacna1e* | calcium channel, voltage-dependent, R type, alpha 1E subunit | 0.4±0.2 |
| *Cacna1f* | calcium channel, voltage-dependent, alpha 1F subunit | 0.1±0.0 |
| *Cacna1g* | calcium channel, voltage-dependent, T type, alpha 1G subunit | 0.0 |
| *Cacna1h* | calcium channel, voltage-dependent, T type, alpha 1H subunit | 64.3±2.2 |
| *Cacna1i* | calcium channel, voltage-dependent, alpha 1I subunit | 0.4±0.3 |
| *Cacna1s* | calcium channel, voltage-dependent, L type, alpha 1S subunit | 0.1±0.0 |
| *Grin1* | glutamate receptor, ionotropic, NMDA1 (zeta 1) | 1.7±0.4 |
| *Grin2a* | glutamate receptor, ionotropic, NMDA2A (epsilon 1) | 0.1±0.0 |
| *Grin2b* | glutamate receptor, ionotropic, NMDA2B (epsilon 2) | 14.2±2.1 |
| *Grin2c* | glutamate receptor, ionotropic, NMDA2C (epsilon 3) | 24.6±4.7 |
| *Grin2d* | glutamate receptor, ionotropic, NMDA2D (epsilon 4) | 2.5±0.1 |
| *Grin3a* | glutamate receptor ionotropic, NMDA3A | 0.7±0.1 |
| *Grin3b* | glutamate receptor, ionotropic, NMDA3B | 0.9±0.2 |
| *Orai1* | ORAI calcium release-activated calcium modulator 1 | 10.4±0.3 |
| *Orai2* | ORAI calcium release-activated calcium modulator 2 | 0.6±0.3 |
| *Orai3* | ORAI calcium release-activated calcium modulator 3 | 15.6±0.0 |
| *P2rx4* | purinergic receptor P2X, ligand-gated ion channel 4 | 29.7±1.4 |
| *Panx1* | pannexin 1 | 26.4±3.4 |
| *Slc24a1* | solute carrier family 24 (sodium/potassium/calcium exchanger), member 1 | 0.1±0.1 |
| *Slc24a2* | solute carrier family 24 (sodium/potassium/calcium exchanger), member 2 | 0.1±0.0 |
| *Slc24a3* | solute carrier family 24 (sodium/potassium/calcium exchanger), member 3 | 2.2±0.2 |
| *Slc24a4* | solute carrier family 24 (sodium/potassium/calcium exchanger), member 4 | 9.9±0.8 |
| *Slc24a5* | solute carrier family 24, member 5 | 7.1±2.1 |
| *Slc8a1* | solute carrier family 8 (sodium/calcium exchanger), member 1 | 12.8±0.4 |
| *Slc8a2* | solute carrier family 8 (sodium/calcium exchanger), member 2 | 0.2±0.0 |
| *Slc8a3* | solute carrier family 8 (sodium/calcium exchanger), member 3 | 0.5±0.0 |
| *Slc8b1* | solute carrier family 8 (sodium/lithium/calcium exchanger), member B1 | 19.3±2.7 |
| *Tmc1* | transmembrane channel-like gene family 1 | 0.2±0.1 |
| *Tmc2* | transmembrane channel-like gene family 2 | 0.0 |
| *Trpa1* | transient receptor potential cation channel, subfamily A, member 1 | 0.3±0.2 |

| | | |
|---|---|---|
| *Trpc1* | transient receptor potential cation channel, subfamily C, member 1 | 0.6±0.4 |
| *Trpc2* | transient receptor potential cation channel, subfamily C, member 2 | 0.0 |
| *Trpc3* | transient receptor potential cation channel, subfamily C, member 3 | 5.7±0.3 |
| *Trpc4* | transient receptor potential cation channel, subfamily C, member 4 | 0.4±0.0 |
| *Trpc5* | transient receptor potential cation channel, subfamily C, member 5 | 4.1±0.4 |
| *Trpc6* | transient receptor potential cation channel, subfamily C, member 6 | 21.5±1.3 |
| *Trpc7* | transient receptor potential cation channel, subfamily C, member 7 | 0.0 |
| *Trpm1* | transient receptor potential cation channel, subfamily M, member 1 | 0.0 |
| *Trpm2* | transient receptor potential cation channel, subfamily M, member 2 | 0.2±0.2 |
| *Trpm6* | transient receptor potential cation channel, subfamily M, member 6 | 2.4±1.2 |
| *Trpm7* | transient receptor potential cation channel, subfamily M, member 7 | 120.0±4.3 |
| *Trpm8* | transient receptor potential cation channel, subfamily M, member 8 | 0.1±0.0 |
| *Trpv1* | transient receptor potential cation channel, subfamily V, member 1 | 0.0 |
| *Trpv2* | transient receptor potential cation channel, subfamily V, member 2 | 1.1±0.5 |
| *Trpv3* | transient receptor potential cation channel, subfamily V, member 3 | 0.3±0.0 |
| *Trpv4* | transient receptor potential cation channel, subfamily V, member 4 | 22.4±2.2 |
| *Trpv5* | transient receptor potential cation channel, subfamily V, member 5 | 0.0 |
| *Trpv6* | transient receptor potential cation channel, subfamily V, member 6 | 0.5±0.1 |

**A**

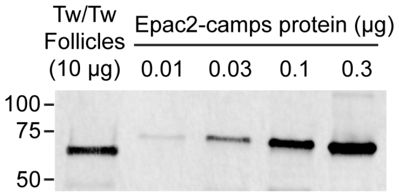

**B**

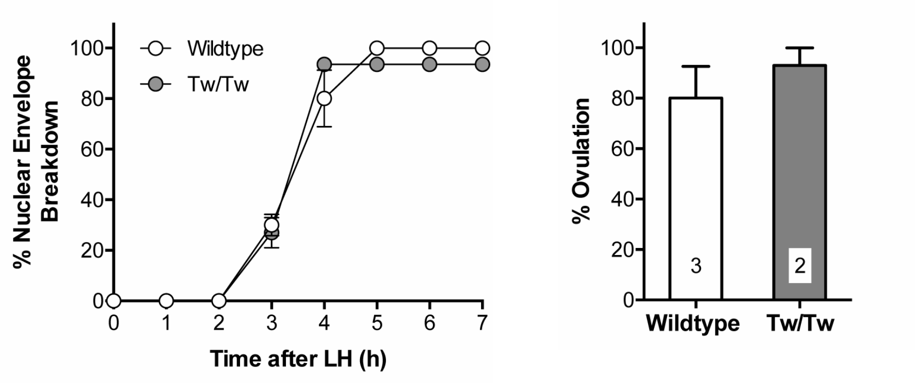

**Figure S1.** Estimate of Twitch-2B concentration in follicles, and normal LH responses of follicles from Twitch-2B homozygous mice. **(A)** Western blot to estimate the concentration of Twitch-2B protein in follicles (320-360 *μ*m diameter) from homozygous (Tw/Tw) mice. 10 μg of follicle protein was separated by SDS-PAGE, in parallel with various amounts of a related YFP and CFP-containing protein (Epac2-camps, >90% pure) that was used as a standard. **(B)** Ovarian follicles from homozygous (Tw/Tw) mice expressing the Twitch-2B sensor show a normal time course of NEBD (left) and a normal percentage of ovulation (right) in response to 10 nM LH. For (A,B), follicles were incubated on Millicell membranes for ~24 hours with 1 nM FSH before use. Data in both panels represent the mean ± s.e.m. of 2-3 experiments with 12-19 follicles each.

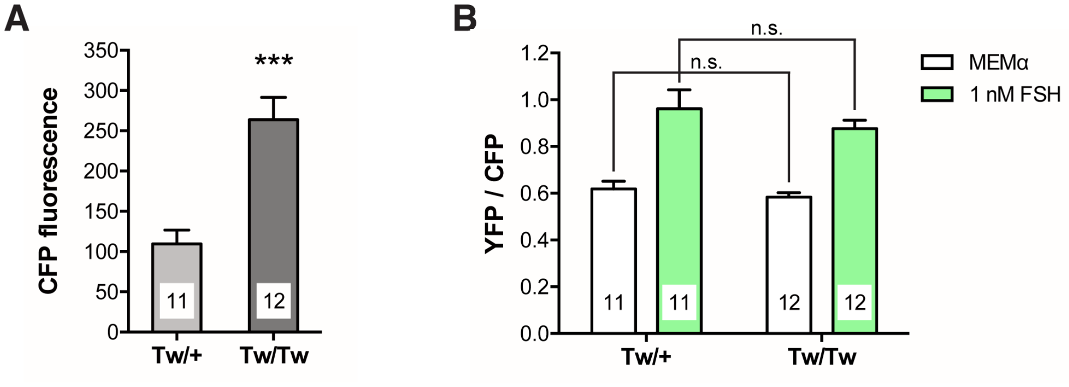

**Figure S2:** Follicles from Twitch-2B homozygous mice express approximately twice as much Twitch-2B sensor as heterozygotes, but both genotypes show similar increases in YFP/CFP ratio in response 1 nM FSH. **(A)** Follicles from mice homozygous for Twitch-2B are significantly brighter than those from heterozygous mice. CFP fluorescence values obtained during the baseline (MEMα) recording period were normalized to the laser power applied. **(B)** Peak YFP/CFP ratios before or after perfusion of 1 nM FSH are not different between follicles from heterozygous and homozygous Twitch-2B mice. Data for (A,B) are from the same follicles as 1 nM FSH bars in Fig. 1D. Numbers within the bars indicate the number of follicles tested. *** indicates $p < 0.001$ by unpaired t-test; n.s. indicates $p > 0.05$ by unpaired t-test. All values represent mean ± s.e.m.

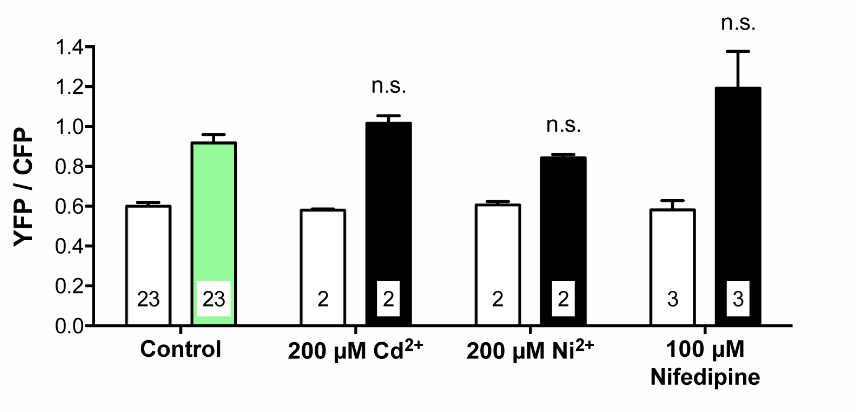

**Figure S3:** The FSH-induced $Ca^{2+}$ increase is not blocked by several inhibitors of voltage-gated $Ca^{2+}$ channels. Peak YFP/CFP ratios before and after perfusion of 1 nM FSH in the presence of 200 $\mu$M $Cd^{2+}$, 200 $\mu$M $Ni^{2+}$, or 100 $\mu$M nifedipine. Open bars indicate the peak YFP/CFP ratio during MEMα perfusion; filled bars indicate peak YFP/CFP ratio after FSH perfusion. For all treatment groups, the peak ratios following FSH perfusion were compared to the control (green) bar by unpaired t-tests; n.s. indicates $p > 0.05$. All values represent mean ± s.e.m.

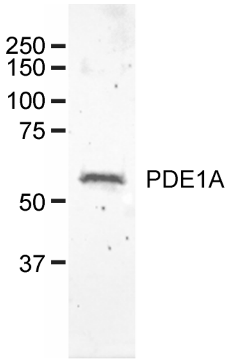

**Figure S4.** Western blot showing the presence of PDE1A in mouse granulosa cells. 2 μg of granulosa cell protein from B6SJLF1/J mice was separated by SDS-PAGE and immunoblotted for PDE1A. Mice were injected at 22 days old with 5 IU equine chorionic gonadotropin. 46 hours later, the ovaries were removed and large antral follicles were punctured with needles to release granulosa cells.

**SUPPLEMENTARY MOVIE LEGENDS**

**Movie S1**. FSH-induced $Ca^{2+}$ increase in a follicle expressing GCaMP6s. The movie shows the time series of images of the follicle region shown in Fig. 2A during sequential perfusion with MEMα (Control) and 1 nM FSH. Images were taken every 10 s, and are displayed at 10 frames per second (fps). Note that the theca cells outside of the follicle show a transient $Ca^{2+}$ increase in response to the mechanical stimulus during perfusion, independent of FSH. Time series is representative of 4 follicles.

**Movie S2**. LH-induced $Ca^{2+}$ increase in a follicle expressing GCaMP6s. The movie shows the time series of images of the follicle region shown in Fig. 5A during sequential perfusion with MEMα (Control), 10 nM LH, and 300 nM LH. Images were taken every 10 s, and are displayed at 10 fps. Three flat-shaped theca cells are seen adhering to the follicle and are brighter than the granulosa cells, indicating a higher level of Twitch-2B expression. These cells also showed $Ca^{2+}$ oscillations in response to LH. Time series is representative of 4 follicles.

**Movie S3**. LH-induced $Ca^{2+}$ increase in a follicle expressing GCaMP6s. The movie shows the time series of images of another representative follicle with a larger field of view, perfused sequentially with MEMα (Control), 10 nM LH, and 300 nM LH. Images were taken every 10 s, and are displayed at 10 fps. A layer of flat theca cells is present, and as seen in Movie S2, showed $Ca^{2+}$ oscillations in response to LH.

**Movie S4**. $Ca^{2+}$ oscillations persist 2 hours after treatment with 10 nM LH in the granulosa cells of a follicle expressing GCaMP6s. Images were taken every 10 s, and are displayed at 10 fps. Time series is representative of 2 follicles**.**

**Movie S5**. $Ca^{2+}$ oscillations persist 6 hours after treatment with 10 nM LH in the granulosa cells of a follicle expressing GCaMP6s. The movie shows the time series of the follicle shown in Fig. 5C. Images were taken every 10 s, and are displayed at 10 fps. Time series is representative of 3 follicles**.**

**Movie S6**. Inhibition of the LH-induced $Ca^{2+}$ increase by inhibition of $G_q$-family G-proteins in a follicle expressing GCaMP6s. The movie shows the time series of images of the follicle region shown in Fig. 6A. The follicle was preincubated for 1 hour with 10 $\mu$M YM-254890 ($G_q$-family G-protein inhibitor), then perfused sequentially with 10 $\mu$M YM-254890, 10 $\mu$M YM-254890 + 10 nM LH, and 10 $\mu$M YM-254890 + 300 nM LH. Images were taken every 10 s, and are displayed at 10 fps. Time series is representative of 3 follicles.
